# Seeing the forest through the trees: Identifying functional interactions from Hi-C

**DOI:** 10.1101/2020.11.29.402420

**Authors:** Ning Liu, Wai Yee Low, Hamid Alinejad-Rokny, Stephen Pederson, Timothy Sadlon, Simon Barry, James Breen

## Abstract

Eukaryotic genomes are highly organised within the nucleus of a cell, allowing widely dispersed regulatory elements such as enhancers to interact with gene promoters through physical contacts in three-dimensional space. Recent chromosome conformation capture methodologies such as Hi-C have enabled the analysis of interacting regions of the genome providing a valuable insight into the three-dimensional organisation of the chromatin in the nucleus, including chromosome compartmentalisation and gene expression. Complicating the analysis of Hi-C data however is the massive amount of identified interactions, many of which do not directly drive gene function, thus hindering the identification of potentially biologically functional 3D interactions. In this review, we collate and examine the downstream analysis of Hi-C data with particular focus on methods that identify significant functional interactions. We classify three groups of approaches; structurally-associated domain discovery methods e.g. topologically-associated domains and compartments, detection of statistically significant interactions via background models, and the use of epigenomic data integration to identify functional interactions. Careful use of these three approaches is crucial to successfully identifying functional interactions within the genome.

## Introduction

The three-dimensional (3D) architecture of the eukaryotic genome has recently been shown to be an important factor in regulating transcription [1–3]. Within the nuclear lamina, DNA is folded into a highly organised structure, allowing transcriptional and regulatory machinery to be in specific nuclear territories for efficient usage. The impact of DNA folding and the resulting physical interactions can have dramatic impacts on the regulation of the genes, enabling non-coding regions such as regulatory elements (e.g. enhancers and silencers) to act on distally located gene promoters with disruption of chromosomal organisation increasingly linked to disease [4–6]. However, while highly organised, the folding structure of the 3D genome can also be highly dynamic to allow for the flexibility and modularity to facilitate regulatory action across a wide-range of cell-types and biological processes, such as development, immune homeostasis, cancer and diseases.

In the recent decades, the development of chromosome conformation capture assays and high-throughput sequencing has facilitated the construction of 3D genome at high resolution, enabling the identification of cell-type and tissue-specific functional interactions between regions in the genome. However, the analysis of such data is complicated by the massive amount of identified physical interactions, hindering the detection and interpretation of interactions that are biologically meaningful. In this review, we introduce the background of 3D genome structure and its components, followed by a summary of the protocols that are commonly used to study 3D genome architecture in recent years, focusing on Hi-C protocols and other derived methods, whilst the use of microscopy to image 3D genome organization has also been recently reviewed [7]. We then thoroughly review current methods for identification of potentially functional interactions and categorise them into three methodological groups.

## Chromosome architecture and gene regulation

Within eukaryotic nuclei, chromosomal DNA is condensed and folded into highly organised 3D structures, with distinct functional domains [8–10]. A key consequence of chromosome folding is that it can bring DNA regions that are far away from each other on the same linear DNA polymer (i.e. intrachromosomal), into close proximity, allowing direct physical contact to be established between regions. Interchromosomal interactions may also play an important role in transcriptional regulation but are less studied. The best characterised examples of this type of interaction include the clustering of ribosomal genes to form the nucleolus and the clustering of olfactory receptor genes to ensure the monogenic and mono-allelic expression in an individual olfactory neuron [11].

The most basic level of chromosome organisation is Chromatin “Loop” structures (Figure 1A). Chromatin loops can either bring distal enhancers and gene promoters into close proximity to increase gene expression, or exclude an enhancer away from the loop to initiate boundaries to repress gene expression [12–14]. The archetypal chromatin looping factor is the CCCTC-binding protein (CTCF) which, along with the Cohesin protein complex, secure the base of a loop and establish the looping structure [3,15–17]. However, other factors such as Yin Yang 1 (YY1) also play an important role in many enhancer-promoter loops [18].

**Figure 1:**
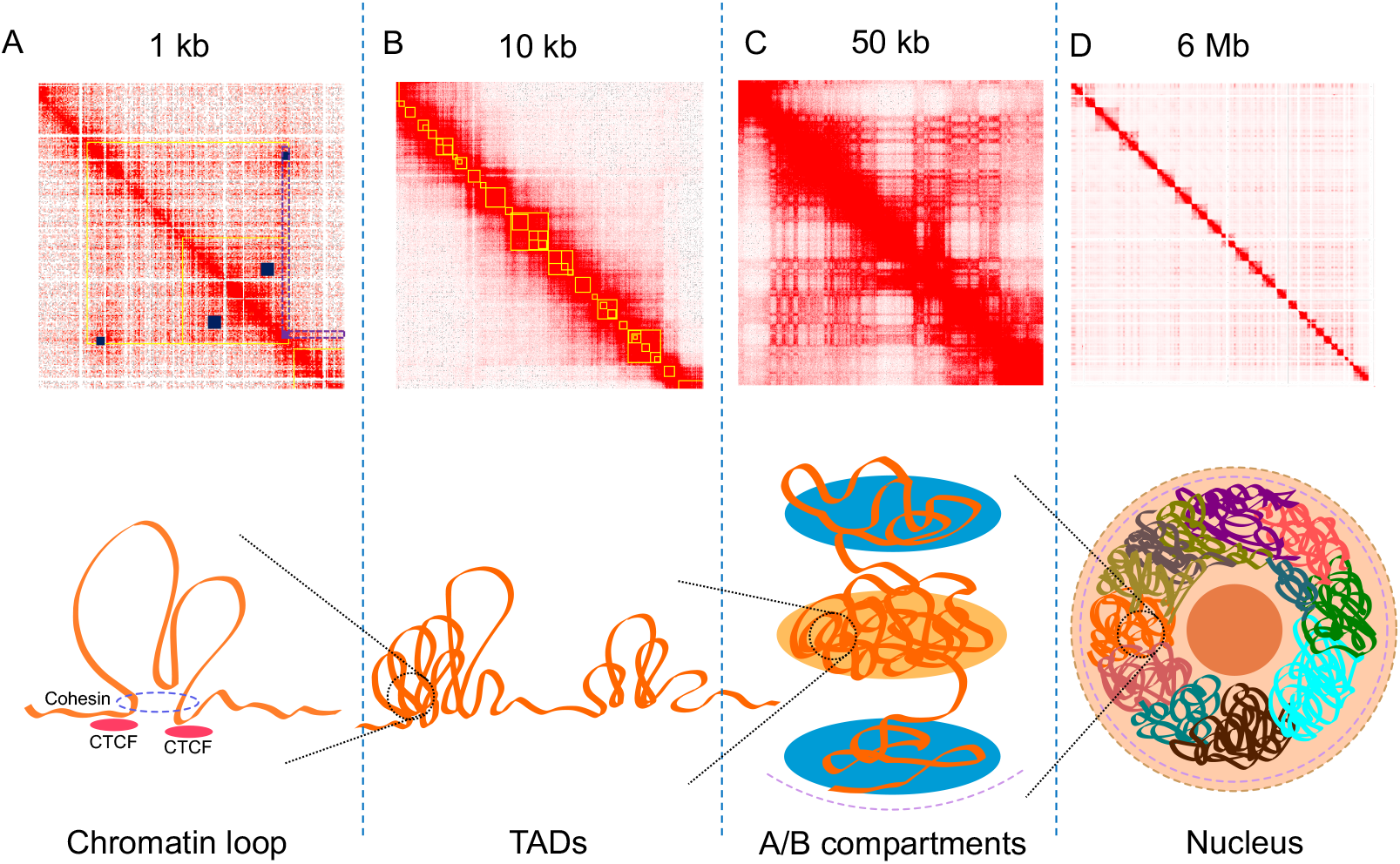
Illustration of genome architecture and the corresponding Hi-C interaction maps. Top panel: interaction heatmaps A, B, C, D are in different scales (kb or Mb per pixel) to correlate with the diagrams of 3D structures in the bottom panel, yellow boxes in A and B are identified TADs and small blue boxes in A indicate chromatin loops. The purple box in A is a frequently interacting region, with its classical “V” shape pattern colored in purple dotted lines. Heatmaps were generated using Juicebox [19] with published Hi-C data of GM12878 [3]. Bottom panel: diagrams of 3D structures in the genome.

Chromatin folding and DNA looping in particular leads to the formation of large scale chromatin structures such as topologically-associated domains (TADs) and chromosome compartments (Figure 1B) [20]. TADs are defined by chromatin interactions occurring more frequently within the TAD boundaries, with TAD boundaries often demarcating a change in interaction frequency [20]. TAD boundaries are also enriched for the insulator-binding protein CTCF and cohesin complex [16,17]. CTCF motif orientation appears to play a role in demarking TAD boundaries with some studies indicating that the majority of identified TADs (~60-90%) have a CTCF motif at both anchor boundaries with convergent orientation [21]. This is consistent with the loop extrusion model, where DNA loops and domains are formed by the extrusion of DNA through the Cohesin complex until the complex encounters convergent CTCF bound at loop anchor sequences [22–25]. Experimental inversion of the CTCF motif orientation has been shown to disrupt the formation of TADs, further emphasizing the important role of the convergent orientation [25]. Recently, a refined definition of TADs was proposed: TADs are formed by extrusion and are strictly confined by boundaries established by ‘architectural’ proteins such as CTCF and Cohesin [26].

The size of TADs can vary from hundreds of kilobases (kb) to 5 megabases (Mb) in mammalian genomes [27,28], and also show significant conservation in related species [29], suggesting that they may serve as the functional base of genome structure and development. Patterns of interactions across regions within a TAD can be further divided into sub-TADs with a median size of 185 Kb using high resolution data [3], enabling finer scale investigation of the genome structure [30,31]. Evidence has shown that TADs are crucial structural units of long-range gene regulation [32,33], with interactions such as promoter-enhancer looping mostly found within the same TADs [34].

At a multi-megabase scale, TADs that are enriched for euchromatin (gene-rich regions) or heterochromatin (gene-poor regions) tend to associate with other TADs of a similar nature leading to the formation of active and inactive domains called ‘Compartments’ (Figure 1C) [35–37]. This compartmentalisation of chromosome folding depicts the global organisation of chromosomes in the nucleus, where compartment A corresponds to gene-dense, euchromatic regions, and compartment B corresponding to gene-poor heterochromatin. Using higher resolution data, the genome can be further grouped into six sub-compartments, compartment A is separated into A1 and A2 whereas compartment B is separated into B1, B2, B3 and B4, with each one associated with specific histone marks [3]. Sub-compartments A1 and A2 are enriched with active genes and the activating histone marks H3K35me3, H3K36me3, H3K27ac and H3K4me1. Sub-compartments A1 and A2 are also depleted in nuclear lamina and nucleolus-associated domains (NADs). B1 domains correlate with H3K27me3 positively and H3K36me3 negatively, B2 and B3 are enriched in nuclear lamina but B3 is depleted in NADs, and B4 is a 11 Mb region, containing lots of KRAB-ZNF genes [3].

The interaction of transcription factors bound at regulatory elements, such as promoters, enhancers and super-enhancers, mediate the transcription level of a gene via interactions which are the direct result of the 3D chromosome structure, but which appear to be long-distance interactions when viewed through lens of a linear chromosome [38–40]. One early and well-characterized example is the gene-gene interaction of the *Fab7* gene in *Drosophila melanogaster.* Using 3D Fluorescence *in situ* Hybridisation (3D-FISH), it was shown that the repression of the gene was regulated by the interaction between a transgenic *Fab7*, which was inserted onto X chromosome, and the native *Fab7* on chromosome 3 [41]. *Hox* gene clusters, essential for patterning the vertebrate body axis, are also governed by a rich enhancer interaction network. Using chromatin conformation capture methods, a number of studies found that the transcriptional activation or inactivation of *Hox* clusters requires a bimodal transition between active and inactive chromatin [20,42–44]. Taken together, the 3D genome structure governing long-distance contacts can build complex gene regulatory networks, allowing for either multiple enhancers to interact with a single promoter or a single enhancer to contact multiple promoters [45]. Disruption of these long range regulatory networks is increasingly being linked to both monogenic and complex diseases [46,47].

## Hi-C assays to quantify chromatin interactions

In order to investigate the 3D genome architecture, a series of protocols called chromosome conformation capture (3C) assays have been developed that specifically capture the physical interactions between regions of DNA [1,2,48–50]. 3C assays work through cross-linking DNA molecules in close proximity via a formaldehyde treatment, preserving the 3D interaction between two genomic regions. The cross-linked DNA is then usually fragmented using a restriction enzyme, such as the 6bp recognition enzyme *HindIII* [20,51] or 4bp cutter *Sau3AI*, and the resultant DNA, ends held in close spatial proximity by the DNA cross-links, are ligated into chimeric DNA fragments. Subsequent steps convert these chimeric DNA fragments into linear fragments to which sequencing adapters are added to create a Hi-C library. The library is then sequenced using high-throughput sequencing technology, specifically limited to Illumina paired-end (as opposed to single-end/fragment) DNA sequencing to enable the accurate identification of the two ends of the hybrid molecule [2].

A suite of 3C-derived high throughput DNA sequencing assays have been developed, including circular chromosome conformation capture sequencing (4C-seq) [48,52], chromosome conformation capture carbon copy (5C) [49], chromatin interaction analysis by paired-end tag sequencing (ChIA-PET) [50] and higher-resolution chromosome conformation capture sequencing (Hi-C) [2], which vary in complexity or the scale of the interactions that are captured. The initial 3C method used PCR to quantify specific ligation products between a target sequence and a small number of defined regions [1]. 4C-seq, known as the “one vs many” method, uses an inverse PCR approach to converts all chimeric molecules associated with a specific region of interest generated in the proximity ligation step into a high throughput DNA sequencing library [52]. 5C increased the number of regions that could be captured by multiplexing PCR reactions [49], albeit developed using older technology (i.e. microarray and 454 sequencing), this approach is considered as the first “many vs many” approach and has been used to examine up to examine the long range interactions of between transcription start sites and approximately 1% of the human genome [53]. ChIA-PET implements a similar approach, however uses a specific, bound protein, generally a transcription factor protein, generating a protein-centric interaction profile [21].

Compared to other approaches, Hi-C is the first “all vs all” method of genome-wide, 3C-derived assay to capture all interactions in the nucleus, allowing for a more complete snapshot of nuclear conformation at the global level [27]. In the initial development of Hi-C, the identification of Hi-C interactions was impacted by the number of spurious ligation products generated as a result of the ligation step being carried in solution allowing for greater freedom for random inter-complex ligation reactions to occur. The resolution of Hi-C interactions in these earlier approaches was also limited by the cutting frequency of a 6-base restriction enzyme, such as *HindIII [2,20,54–56]*. To address these issues, an *in situ* Hi-C protocol was developed [3], where the ligation steps were performed within the constrained space of the nuclei, reducing the chance of random ligation [57]. Furthermore, *in situ* Hi-C used a 4-base-cutter (such as *MboI*) for digestion, increasing the cutting frequency in the genome and improving the resolution of captured interactions [3]. Using this method, the first 3D map of the human genome was constructed using the GM12878 cell line with approximately 4.9 billion interactions [3], enabling interaction resolution at the kilobase level. In recent years, the *in situ* Hi-C protocol has been developed further to target different technical and/or biological questions (Table 1).

**Table 1:** Different Hi-C-derived methods. Optimizations indicate their modification in their protocols compared to traditional Hi-C.

| Hi-C flavours | Optimizations | Advantages | Reference |
| --- | --- | --- | --- |
| Traditional Hi-C | - | - | [2] |
| in situ Hi-C | Nuclear ligation; 4-based cutter | Allowing higher resolution data generation | [3] |
| DNase Hi-C | DNase I to digest crosslinked DNA | Improving capture efficiency, reducing digestion bias | [58] |
| Micro-C | Micrococcal nuclease to digest crosslinked DNA | Improving capture efficiency, reducing digestion bias | [59] |
| Capture HiC | RNA baits to subset specific chromatin contacts | Reduce sequencing cost, focus on a subset of interactions | [60] |
| HiChIP | Chromatin Immunoprecipitation (ChIP) to subset bound chromatin contacts | Reduce sequencing cost, focus on a subset of interactions | [61] |
| scHiC | Single cell technology | Allowing investigation of 3D variation at single cell level | [62] |
| scMethyl-HiC/ sn-m3C-seq | Single cell methods using bisulfite conversion | Allowing jointly profiling of DNA methylation and 3D genome structure in single cell level | [63], [64] |

Owing to the vast complexity of the Hi-C ligation products generated, it is often too costly to sequence samples to a sufficient depth to achieve the resolution necessary to investigate specific interactions such as promoter-enhancer interactions, leading to the development of capture Hi-C-derived methods (CHi-C) [60]. CHi-C employs a sequence capture approach, using pools of probes complementary to thousands of restriction fragments, to enrich for molecules containing the region of interest from the HiC library. This significantly reduces the complexity of the libraries and enables a significant increase in the number of detectable interactions within specific regions without the need for ultra-deep sequencing. Therefore CHi-C, has been used in many cases to analyse specific types of long-range interactions, such as interactions linked to promoter or enhancer regions. For example, CHi-C was recently used to characterise promoter interactions in 17 human primary hematopoietic cells to demonstrate the highly cell-type specific nature of many promoter interactions even with a group of related cell types [39].

Similar to many other high-throughput sequencing approaches, Hi-C continues to be modified to improve the efficiency and resolution of the approach. DNase Hi-C was developed to address the bias introduced through the use of restriction enzymes (e.g. *MboI* recognizes GATC), due to the uneven distribution of restriction sites throughout the genome [58,65]. Instead, DNase Hi-C replaces the restriction enzyme digestion of cross-linked DNA with the endonuclease DNase I that has a much reduced DNA sequence specificity to reduce bias in identifying Hi-C interactions. Commercial Hi-C library preparation kit such as Omni-C kit from Dovetail Genomics [66] exploits the use of DNase and is designed specifically to overcome limitations of only capturing Hi-C interactions near restriction sites. Similar to DNase Hi-C, Micro-C uses micrococcal nuclease digestion, enabling the resolution at a 200 bp to ~4 kb scale in budding yeast [59].

The integration of Hi-C with other genomic applications, such as chromatin immunoprecipitation (ChIP), bisulfite treatment or single cell technologies has also occurred. The ChIP approach, HiChIP i.e. Hi-C with an additional ChIP step, has been used to enrich a Hi-C library for interactions associated with specific bound proteins [61], increasing the resolution of the library while reducing the sequencing cost. While similar to ChIA-PET, HiChIP generates similar numbers of interactions while requiring significantly less starting material due to the order in which the proximity ligation and chromatin immunoprecipitation are performed. In HiCHIP, the chromatin immunoprecipitation step occurs following the proximity ligation of fragmented cross-linked DNA maximising the yield of the inefficient chromatin immunoprecipitation step. In contrast in ChIA-PET the chromatin immunoprecipitation step is performed prior to ligation resulting in a loss in material [61]. Single cell Hi-C (scHiC) combines the Hi-C protocol with single cell technologies, providing the ability to investigate 3D genome structure variation at a cell-to-cell level [62,67–70]. Furthermore, another method called sn-m3C-seq adapted single cell and bisulfite conversion methods with Hi-C to allow robustly distinguish cell types in single cell level [64]. Methyl-HiC and scMethyl-HiC have also been developed to jointly profile the DNA methylation and 3D genome structure on both bulk and single cell level [63]. Recent studies have revealed that DNA methylation is able to impact 3D genome structure via polycomb complexes, which play an important part in respressing key developmental genes [71].

As the development of Hi-C approaches continue, it is essential that computational methods are standardized in order to provide consistent results that are comparable across species or cell-types. In the next section, we review the current data processing methods that are used in standard Hi-C sequencing approaches.

## Prioritisation of chromatin interactions

Methodologies to extract meaningful, functional information from the massive number of interactions identified through Hi-C data can be categorized into three groups: structural-based methods, detection of significant interactions and data integration (Figure 2). The first approach is to define structures such as A/B compartments and TADs, based on the 2D interaction patterns across the genome. The second approach is to investigate only a subset of Hi-C interactions, that are identified from a statistical test based on a trained model. Finally, taking advantage of the publicly available databases, the third approach is to prioritise interactions that are more likely to be biologically relevant through the investigation of genomic and epigenomic information. These approaches are not mutually exclusive and in many cases can be combined to address specific questions in genome organisation and gene regulation.

**Figure 2:**
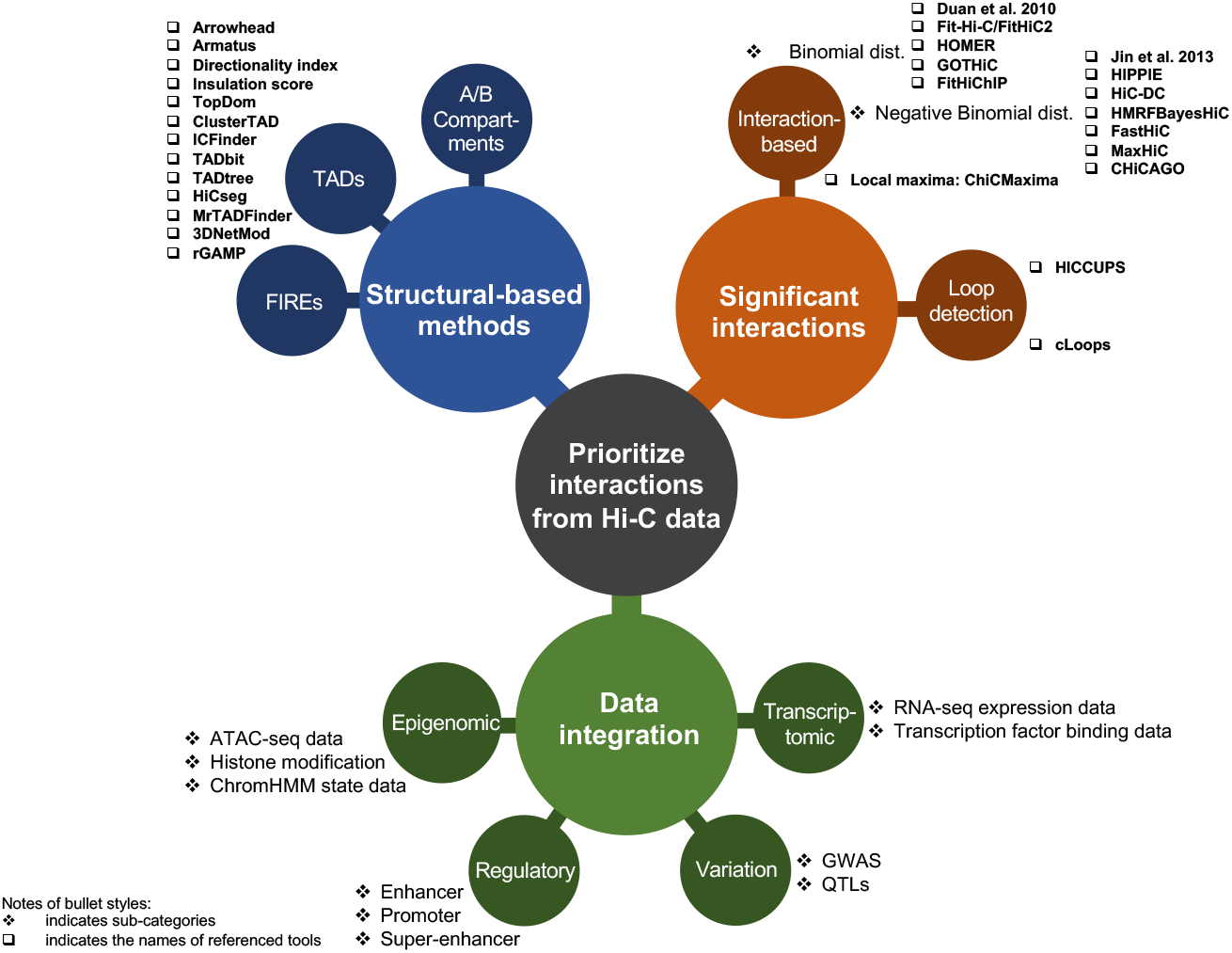
Approaches to prioritise interactions from Hi-C datasets. In this review, we categorised the approaches to identify functional interactions into three ways, including significant interactions identification, structures summarisation and data integration. Referenced tools and sub-categorical analyses are marked on the figure with boxes and stars respectively.

## Structural-based identification methods

Methods that identify structural aspects of chromatin interactions (i.e. A/B compartments and TADs) are employed as an avenue to reduce the dimensionality of the 3D interaction profile by clustering or summarising regions with similar patterns across the genome. The A/B compartments are commonly predicted with normalised Hi-C matrices generated using Valina Coverage (VC) [2], Knight and Ruiz’s method (KR) [72] or iterative correction and eigenvector decomposition (ICE) [73]. Normalised data is then used to calculate Pearson’s correlation and through principal component analysis (PCA), the eigenvectors of the first (or second) principal component (PC) are usually used to assign bins to A or B compartments. Current analysis toolkits, such as Juicer [74] and FAN-C [75], have optimised correlation matrix functions to identify A/B compartments from Hi-C matrices without significant taxes on memory and computational resources.

As detailed above, TADs are defined as structures with interactions that occur within TADs rather than across TADs [20]. As such, they are often identified by detecting boundary characteristics where the interaction signals can be observed to be significantly different at both sides [20,76]. Currently, there are over 20 commonly used TADs callers that have been developed using various methodologies. For instance, arrowhead [3], armatus [77], directionality index [20], insulation score [76] and TopDom [78] use their own linear scoring system, clusterTAD [79] and ICFinder [80] are based on clustering, TADbit [81], TADtree [82] and HiCseg [83] use statistical models; and MrTADFinder [84] and 3DNetMod [85] rely on network-modelling approaches [28,86]. Although comparisons reveal low reproducibility among tools, especially in the number and mean size of identified TADs, recent reviews [28,86] have suggested a preference for TAD callers that allow for the detection of nested TADs or overlapped TADs, such as rGMAP [87], armatus, arrowhead and TADtree.

While theoretically similar to TAD calling, frequently interacting regions (FIREs) are also commonly used to describe structural interaction characteristics. Defined as genomic regions with significant interaction profile, FIREs exhibit strong connectivity with multiple regions in the chromosome neighbourhood [55]. FIREs can be easily visualised on the Hi-C interaction map, with interacting signals appearing from both sides of the FIREs, forming a characteristic “V” shape (Figure 1A). FIREs are typically less than 200 Kb in size, hence are regarded as local interaction hotspots [55,88]. Unlike TADs and compartments, which exhibit a certain level of conservation across cell types (about 50~60% and 40%, respectively) [3,20,55,89], FIREs appear to be cell type- and tissue-specific and are often located near key cell phenotype-defining genes. However, similar to TADs, FIREs formation seems to be dependent on the Cohesin complex, as its depletion results in decreasing interactions at FIREs [55]. They are also enriched for super-enhancers, suggesting FIREs play an important role in the dynamic gene regulation network [90,91].

## Methods for identification of significant chromatin interactions

In order to prioritize potentially meaningful chromatin interactions, statistical significance is assigned to Hi-C interactions by comparing them to a background model and assessing the probability of observing the experimental set of counts if the background model were the underlying method of generating observed counts. The interaction frequency generally decays with increasing linear distance, and by applying this background model meaningful interactions can be identified through a higher than normal frequency. Here we summarize the current methodologies of significant interactions identification and categorised them into two groups; interaction-based methods, which define a background signal model by considering the read count of any pair of interactions, and loop-based methods, which account for interactions in the neighbouring areas to identify peak interactions with statistical significance.

### Interaction-based methods

Initial studies assigning statistical significance to Hi-C interactions separated interactions into intra-chromosomal interactions (within the same chromosome) and inter-chromosomal interactions (across two chromosomes), followed by a binomial distribution to assign confidence estimates for inter-chromosomal interactions [92]. A binning method is then used to account for the characteristic pattern of intra-chromosomal interactions, with the observed interacting probability decaying as the genomic distance increases linearly. This is then used to compute interacting probabilities for each bin separately and assigning statistical significance using the same binomial distribution as used for inter-chromosomal interactions [92]. Based on the same binomial distribution concept, Fit-Hi-C uses spline fitting procedure instead of binning, reducing the bias of artifactual stair-step pattern [93]. Additionally, Fit-Hi-C also incorporates an extra refinement step using a conservative model with stringent parameters to remove outlier interactions, which can be applied iteratively, to achieve a more accurate empirical null model. However, Fit-Hi-C was initially limited by only allowing bin sizes larger than 5 kb to compute significance due to the heavy memory usage when dealing with higher-resolution data. However this has been improved with recent updates [94], and is now able to handle data with high resolution (bin sizes from 1 to 5 kb). Other new features include accepting multiple input formats so that it is compatible with different Hi-C analysis pipelines are acquired in the latest version of Fit-Hi-C. Another similar tool is included in the Homer toolkit [95], which accounts for biases such as sequencing depths, linear distance between regions, GC bias and chromatin compaction to establish a background model to estimate the expected interaction count between any two regions, followed by the use of a cumulative binomial distribution to assign significance to interactions. GOTHiC [96] also uses relative coverage of two interacting regions to estimate both known and unknown biases, followed by a cumulative binomial distribution to build the background model to identify significant interactions

The Negative Binomial distribution is commonly utilised in the analysis of count-based data, including popular RNA-seq analysis tools such as edgeR [97] and DEseq2 [98], and has been implemented in a number of Hi-C programs such as HIPPIE [54,99]. This method uses a negative binomial model to estimate the statistical significance of the interactions in one fragment region (< 2 Mb) while accounting for restriction fragment length bias and interacting probability distance bias simultaneously. However, negative binomial models can be confounded by many bins with zero counts [100] and a number of programs have developed approaches to account for “zero-inflated” observations. HiC-DC, for example uses a hurdle negative binomial regression model to identify significant interactions [100], modelling the probability of non-zero counts and the rate of observed counts as separate components of the model.

While physical interactions between loci found in close linear proximity are likely to be more prevalent in Hi-C datasets, a known bias in Hi-C libraries is the correlation between two nearby restriction fragments brought about by ligation events. Ligation events can be the result of bias or random collision events between restriction fragments during library preparation, so with high coverage sequencing, false signals can impact the identification of significant interactions [54]. To tackle this problem, HMRFBayesHiC uses a negative binomial distribution to model observed interactions [54], followed by a hidden Markov random field model to account for the correlation between restriction fragments, and to model interaction probabilities [101]. This implementation required significant resources to run, leading to the development of FastHiC [102], which enables higher accuracy of interaction identification and faster performance. Recently, another tool called MaxHiC also based on negative binomial distribution was developed [103]. Compared to other tools, all parameters of the background model in MaxHiC are established by using the ADAM algorithm [104] to maximize the logarithm of likelihood of the observed Hi-C interactions. Significant interactions identified by MaxHiC were shown to outperform tools such as Fit-Hi-C/FitHiC2 and GOTHiC in identifying significant interactions enriched between known regulatory regions [103].

Compared to traditional Hi-C protocols, Capture Hi-C (CHi-C) requires different analytic methods due to the extra bias driven by the enrichment step in the protocol. Capture libraries can be regarded as a subset of the original Hi-C library, meaning the interaction matrix of CHi-C is asymmetric, and interestingly not accounted for in traditional normalisation methods [60,105]. Because of this, many analysis approaches are specifically designed for CHi-C data analysis. CHiCAGO (Capture Hi-C Analysis of Genomic Organisation) was developed to account for biases from the CHi-C protocol and identify significant interactions [105], using a negative binomial distribution to model the background local profile and an additional Poisson random variable to model technical artefacts [105]. CHiCAGO uses the implicit normalization method ICE [73] and multiple testing stages based on p-value weighting [106] to carefully identify significant interactions from each CHi-C dataset [105]. Another CHi-C-specific tool called ChiCMaxima was developed to identify significant interactions by defining them as local maxima after using loess smoothing on bait-specific interactions [107]. Compared to CHiCAGO, ChiCMaxima’s approach is more stringent and exhibits a more robust performance when comparing biological replicates datasets [107]. As well as being applicable to conventional HiC data, MaxHiC is also able to identify significant interactions in CHi-C data [103] and offers robust performance to identify regulatory areas compared to CHi-C-specific tools including CHiCAGO [103].

Like the other capture approaches, HiChIP cannot use traditional (Hi-C-specific) interaction callers (e.g. Fit-Hi-C or GOTHiC) due to the inherent biases associated with an enrichment with specific immunoprecipitation targets [61]. Hichipper was developed to firstly identify ChIP peaks while accounting for the read density bias in restriction fragments, enabling a more accurate identification of interactions from HiChIP dataset [108]. While hichipper does not implement any function to identify significant interactions, FitHiChIP was developed to account for non-uniform coverage bias and distance bias in restriction fragments using a regression model, together with 1D peak information in a spline fitting procedure to accurately identify significant interactions from HiChIP data [109].

### Chromatin loop detection

Based on the loop extrusion model of the 3D genome [23–25], chromatin looping structures can be regarded as the basic unit of 3D genomic architecture and play an important role in the regulatory process, by bringing distal promoter and enhancer elements together or excluding enhancers from the looping domain [12–14]. Chromatin loops from Hi-C data were first defined by searching for the strongest “pixel” on a normalised Hi-C contact map (Figure 1A). By comparing all pixels in a neighbouring area, the two bins associated with the pixel are regarded as the anchor points of a chromatin loop when the interaction frequency of the pixel is the strongest among the neighbouring pixels [3]. A searching algorithm named Hi-C Computational Unbiased Peak Search (HICCUPS) was developed to rigorously search for these pixels, enabling the identification of chromatin loops from Hi-C data [3]. Somewhat similar to TADs, published information on chromatin loops demonstrates structural conservation between a number of human cell lines (~55-75% similarity), and between human and mouse (about 50% similarity), suggesting conserved loops may serve as a basic functional unit for the genome [3]. However, loop detection using HICCUPS requires high resolution data with extremely high sequencing depth. For example, over 15 billion unique interactions from more than 5 tera bases (TBs) of raw sequencing Hi-C data were required by HICCUPS to identify 10,000 unique loops [3]. The current development of loop detection algorithms have focused on deep learning approaches, such as DeepHiC [110] using generative adversarial networks, as well as HiCPlus [111] and HiCNN [112] which use deep convolutional neural networks. Such methods can potentially be used to increase the resolution of Hi-C to achieve necessary depth to identify chromatin loops.

Hardware requirements to identify loops in high-resolution data is also extremely restrictive with HICCUPS requiring specific architectures (i.e. NVIDIA GPUs) to identify looping patterns. In order to reduce the computational cost, an approach called cLoops was implemented which identifies peak interactions from chromatin contact map [113]. cLoops initiates loop detection by finding candidate loops via an unsupervised clustering algorithm, Density-Based Spatial Clustering of Applications with Noise (DBSCAN) [114], which enables computing statistical significance of interactions with less amount of input and reduced computational resources. Candidate loops are then compared with a permuted background model, based on the interaction decay over linear distance, to estimate statistical significance.

### Functional interaction identification via omics-based data integration

The fundamental motivation for identifying interacting regions across a genome is to establish how non-coding regions of the genome impact gene expression [1,115,116]. However functionally-relevant interactions, whether this be chromatin interactions between gene promoters and enhancers or transcription factor binding mechanisms, are often established in a cell-type specific manner [51,60]. By integrating local or publicly available genomic, transcriptomic and epigenomic datasets, significant interactions (such as the ones identified in the programs above) are able to be prioritised to the functionally relevant domains in which they belong.

For regulatory elements, such as promoters and enhancers, the identification of open or closed chromatin, through assays for DNaseI-hypersensitivity, ATAC-seq or specific histone modifications, is especially valuable in determining the functional potential of an interaction. Public databases such as the FANTOM5 project [117], the NIH Roadmap Epigenomics project [118], the EU Blueprint project [119] and ENCODE [120,121] contain an enormous number of essential regulatory information, often to a single molecular level. For example, the Roadmap Epigenomics project uses a multivariate hidden Markov model to learn the information of multiple histone marker signals and/or DNase I signals to predict discrete chromatin states [114]. The initial analysis consisted of 15 predicted chromatin states across 127 cell-type specific epigenomes, containing promoters-associated states such as Active TSS, Flanking Active TSS, Bivalent/Poised TSS and Flanking Bivalent TSS and enhancer-associated states such as Enhancers, Genic Enhancers and Bivalent Enhancers [118]. Expanding on the initial 15 predicted states the roadmap project now also provides, 18-state models from 98 epigenomes, a 50-state model of 7 deeply-profiled epigenomes, and a 25-states model from 127 epigenomes, which based on imputed data to create finer-grained detail [122]. While chromatin states data are human specific, mouse tissue-specific chromatin states data are available in a public GitHub repository from a recent study [123]. Predicted chromatin states have been widely used as an integrative annotation tool for Hi-C or CHi-C studies, enabling cell type/tissue-specific information across large unannotated regions of the genome [55,124–126].

One crucial annotation element relevant to the identification of functional interactions is the presence of super-enhancers (SEs). SEs are clusters of enhancers exhibiting significantly higher levels of active enhancer marks compared to most enhancer elements within a cell type and are enriched with numerous Transcription Factor-Binding Sites (TFBS) [127]. These regions act as “regulatory hubs”, which are higher-order complexes consisting of interactions between multiple enhancers and promoters at individual alleles [128–130]. The formation of these regulatory hubs are proposed to be the consequence of the high level of TF and co-factor localisation to the SE interacting to form a biomolecular condensate by a phase separation model [131,132] [133–136]. SE can be identified from H3K27ac ChIP-Seq using the ROSE algorithm [137], and currently SE information can be easily accessible from databases such as AnimalTFDB [138], PlantTFDB [139], GTRD [140], SEdb [141], dbSUPER [142] and SEA [143], allowing hubs to be identified at a cell specific level. A number of studies have used Hi-C or other types of chromosome conformation capture data to establish the idea that TFs and SEs are key regulators that can mediate multiple gene expression regulations three-dimensionally, and they are important for cell identity and development [38,137,144–147].

One of most challenging problems in human genetic studies is that most of the identified genetic variation associated with phenotypic traits are located in non-coding regions, with many studies inundated that enhancers and super-enhancers are particularly enriched for disease-associated genetic variation [148,149].Therefore, integration of genetic variation data in the form of aggregated or domain-specific SNP databases with functionally important interactions identified from Hi-C data has been used to investigate disease association. Databases such as GWAS catalog [150], ImmunoBase [151], GWAS Central [152], GWAS ALTAS [153] and GWASdb [154] contain information of the level of genetic association of each variant to specific diseases, which are invaluable data to be integrated in a high-dimensional interaction dataset. When aggregated with expression quantitative trait loci (eQTL) information (i.e. variants that impact expression on genes in proximal or far genes) from databases such as the GTEx consortium [155], interactions can be interpreted by their impact on gene expression. As an example Greenwald *et al.* has recently used pancreatic islet-specific data to investigate the risk gene loci of Type 2 Diabetes (T2D) [156]. In their work they combined gene and enhancers interaction maps generated from Hi-C data, together with variant and gene expression linkage data, provided by tissue-specific eQTL analysis, to establish an enhancer network for T2D risk loci. In support of genetic variation at enhancers influencing transcriptional regulation, Yu *et al.* used HiC data to demonstrate that eQTLs tend to be in close spatial proximity with their target genes [157]. Additionally, by integrating the information of GWAS SNPs, TF binding site prediction and Hi-C/CHi-C data, studies have shown that non-coding SNPs can affect the binding of TFs, and can further potentially affect multiple target genes via 3D interactions [158,159].

## Future prospects

The investigation of 3D chromosome structure can provide novel insights into the complex regulatory network in the genome. The development of Hi-C and its derived protocols have facilitated the studies of the 3D genome structure, generating numerous high quality datasets. However, due to the complexity of the Hi-C library preparation and analysis, the biologically meaningful, small-scale interactions may still lack sufficient signals, hindering the detection and interpretation of 3D interactions. The approaches that we presented in this review all aim to reduce the complexity of 3D interaction data, narrowing down information based on structure, statistical inference and additional lines of experimental evidence (i.e. cell-type specific epigenomic data).

Incremental development of Hi-C calling applications (chromatin loops, TADs etc) has continued with a focus on correcting biases introduced by library preparation and sequencing. As more and more sequencing data is deposited on open-access data repositories such as NCBI Short Read Archive (SRA) [160] and European Nucleotide Archive (ENA) [161], has allowed the development of novel Machine Learning models trained on known interactions to identify novel patterns when applying these models to new datasets. Incorporation of publicly-available cell type/tissue-specific epigenomics data into these machine learning models of chromatin interactions will allow for more accurate predictions on the molecular mechanisms by which diseases-associated genetic acts. In the future, such models of 3D interactions can potentially be used as markers for disease screening and used for personalised medicine development.

Another exciting development has been the development of single cell Hi-C approaches. This allows for a more detailed analysis of the dynamic nature of chromatin interactions than provided by bulk population based Hi-C assays [70,162]. In particular, single-cell approaches offer significant promise in identifying how interactions change during time course-dependent processes, such as cell-type differentiation processes. Single-cell Hi-C analyses have already been successfully conducted [62,68,70,163,164] and algorithms developed that are able to detect interactions despite sparse data matrices and significant difficulties with processing [165,166]. As single-cell Hi-C methods continue to be developed, in the future Hi-C will be able to complement RNA Velocity or “pseudo-time” analyses [167] that aim to track cell fate interactions in real-time.

Although the development in protocol efficiency, parallel algorithmic improvements are likely to improve current approaches for identifying 3D interactions. Additional imaging technologies such as real-time signal Fluorescence *in situ* hybridization (FISH) and advanced imaging approaches such as STORM imaging have been used to visualise the nuclear organization in living cells and leading to the identification of clusters of clutch domains that are thought to correspond to TAD [7,168]. Lastly the ability to engineer specific mutations in DNA through genome editing technology such as the CRISPR-Cas9 system [169,170], means that future experiments using Hi-C and 3D imaging in-parallel with genetically modification of genomes will vastly improve our understanding of how variation may impact genomic structure, and the regulations of gene expression.

## Conclusion

In this review, we first introduced the three-dimensional chromosome architecture in different scales, followed by presenting the chromosome conformation capture assays, with a focus on Hi-C and its variations, which are the state of the art methods for investigating the 3D genome structure. Lastly, we comprehensively reviewed methodologies that are developed to reduce the complexity of 3D physical interactions identified from Hi-C datasets to detect potential functional interactions. We also categorised the methods into three types, including structural-based detection methods, significant chromatin interactions identification methods and data integration methods. Taken together, by utilizing these methods carefully, we are able to detect physical interactions with biological meaning and impact from complicated Hi-C dataset, which may serve a purpose in diagnosis and precision medicine.

## Key points

- Hi-C and its variation assays can be used to investigate 3D chromosome architecture in different scales.
- This review provides an overview of popular methodologies used to identify potential functional physical interactions from Hi-C and Hi-C derived datasets.
- The reviewed methods can be categorised into three groups, including structural-based detection methods, significant chromatin interactions identification methods and data integration methods.

## Funding

This work was supported by a 2017 National Health and Medical Research Council (NHMRC) project grant (*1120543*).

**Ning Liu** holds a BSc in Biotechnology Science and an MSc in Biomedical Science. Currently, he is working at the South Australian Health and Medical Institute (SAHMRI), in Adelaide, Australia, conducting his doctoral studies in the field of Bioinformatics, with **Dr James Breen**.

**Wai Yee Low**, PhD, is a Bioinformatics Data Scientist at the Davies Research Centre, University of Adelaide and he specialized on ruminant genomics. Prior to this he was an Assistant Professor and Head of Products and Services at the Perdana University Centre for Bioinformatics.

**Hamid Alinejad-Rokny**, PhD, is a UNSW Scientia Fellow and leads Systems Biology and Health Data Analytics Group at UNSW Graduate School of Biomedical Engineering. His research focuses on using cutting-edge systems biology and machine learning techniques in conjunction with omics data to understand the mechanisms implicated in genetic diseases and cancers.

**Stephen Pederson**, PhD, is a Post-Doctoral Bioinformatician in the Dame Roma Mitchell Cancer Research Laboratories, Adelaide Medical School, University of Adelaide. Prior to this he ran the University of Adelaide Bioinformatics Hub for 6 years, and has a keen interest in transcriptomics and gene regulation

**Timothy Sadlon**, PhD, is a senior Post-Doctoral fellow in the Molecular Immunology group headed by Professor Barry. He has a keen interest in the role 3D chromatin structure plays in transcriptional regulation in the immune system.

**Simon Barry**, PhD, is a Molecular Immunology group leader, with research interests in the genetic basis of disease and functional annotation of 3D genomic interactions to explain altered phenotype by combining transcriptomics, epigenetics, conformation and accessibility.

**James Breen**, PhD, is the leader of the SAHMRI Bioinformatics Core, with research interests including developing methods for clinical cancer sequencing, and investigating epigenetic and structural gene regulation using multi-omic datasets.

